# A Fast Detection Method for Wheat Mould Based on Biophotons

**DOI:** 10.1101/2020.10.13.337246

**Authors:** Gong Yue-hong, Yang Tie-jun, Liang Yi-tao, Ge Hong-yi, Chen Liang

## Abstract

Mould is a common phenomenon in stored wheat. First, mould will decrease the quality of wheat kernels. Second, the mycotoxins metabolized by mycetes are very harmful for humans. Therefore, the fast and accurate examination of wheat mould is vitally important to evaluating its storage quality and subsequent processing safety. Existing methods for examining wheat mould mainly rely on chemical methods, which always involve complex and long pretreatment processes, and the auxiliary chemical materials used in these methods may pollute our environment. To improve the determination of wheat mould, this paper proposed a type of green and nondestructive determination method based on biophotons. The specific implementation process is as follows: first, the ultra-weak luminescence between healthy and mouldy wheat samples are measured repeatedly by a biophotonic analyser, and then, the approximate entropy and multiscale approximate entropy are separately introduced as the main classification features. Finally, the classification performances have been tested using the support vector machine(SVM). The ROC curve of the newly established classification model shows that the highest recognition rate can reach 93.6%, which shows that our proposed classification model is feasible and promising for detecting wheat mould.

## Introduction

Wheat, as a type of global grain, is one of the staple foods that human beings and animals rely on throughout the world. The history of wheat cultivation can be traced back to ten thousand years ago, and it has become the second most cultivated crop in the world due to its high productivity and strong adaptability [1]. As the world population has increased over the last decade, the consumption of wheat has also increased [2], which can be seen in Fig. 1.

**Fig. 1.**
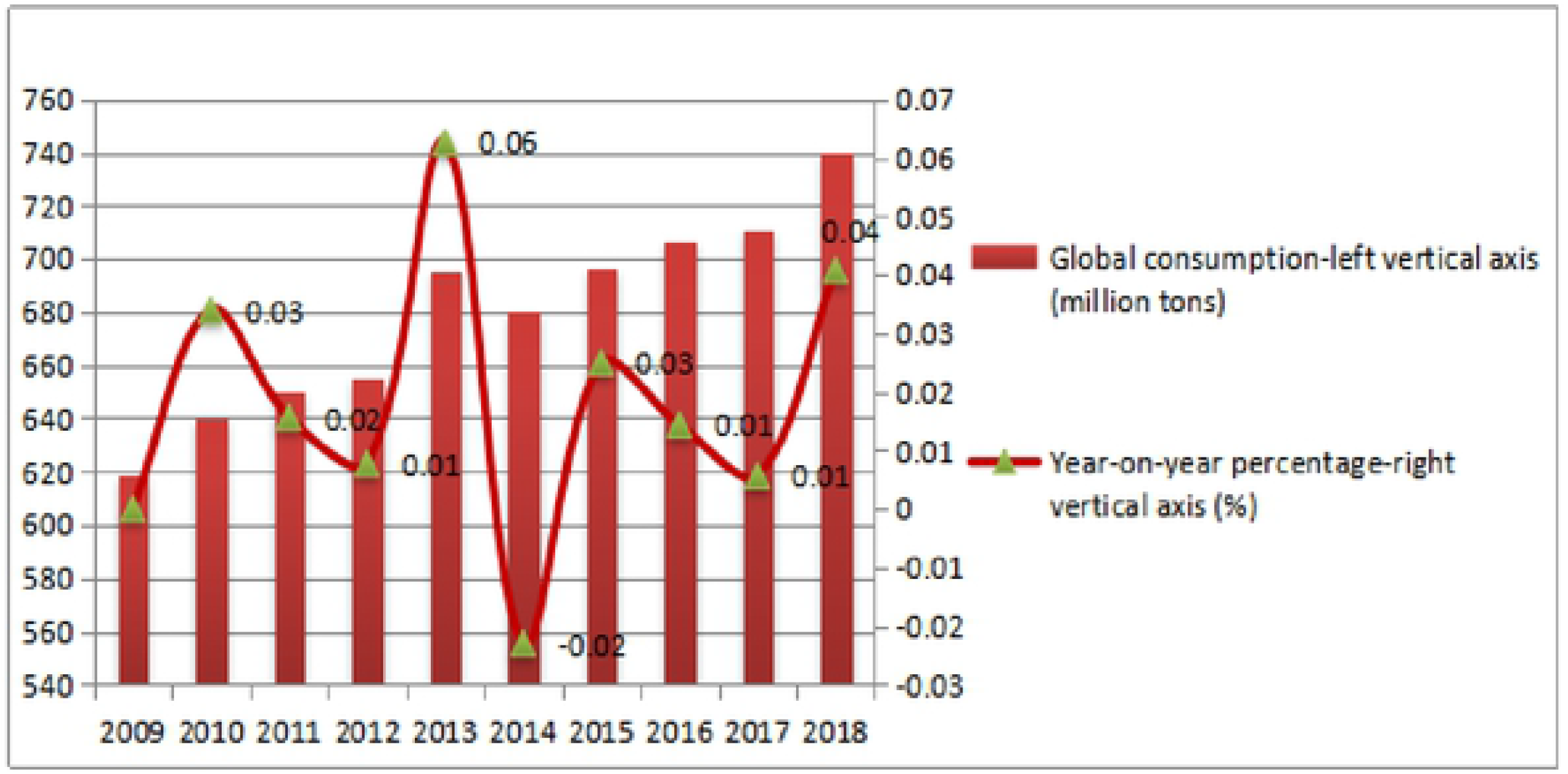
Global Consumption of Wheat and Year-on-Year Percentage from 2009 to 2018.

When a suitable surrounding moisture and temperature is achieved, microorganisms make great contributions to the wheat mould phenomenon, thus affecting the quality and quantity of stored wheat [3]. Since mould is inevitable during the wheat storage period, the health of human beings will be extremely threatened once certain edible food that is made using mouldy wheat as raw materials are available in their daily lives [4]. Many mycotoxins are metabolized by mouldy wheat, among which aflatoxin B1 (AFB1) is the most striking contaminant and has the strongest carcinogenicity [5]. Nearly one quarter of crops in the world are contaminated by aflatoxins before or during their storage period according to the Food and Agriculture Organization (FAO). Once feeds and foods are made of mouldy kernels, the AFB1 carried within will cause a series of illnesses, such as retarded growth, immune suppression, human or animal death, and so on [6]. Therefore, the development of fast and green techniques for detecting AFB1 in stored wheat kernels is very necessary to ensure human and animal safety.

The study of biological photons can be traced to 1923 when the Russian biologist Gurwitsch used biological detectors to test the roots of onions and found a special phenomenon: onion cells can produce faint light that can stimulate other cells to accelerate their cell division [7]. The Italian scientist Coli placed some plant buds on detectors with photomultiplier tubes for measurement and observed an ultra-weak light emission phenomenon [8]. In the later 1970s, led by West German physicians, biophoton research conducted many experiments and made striking progress [9]. In the 1980s, biophoton technology was applied to spontaneously detect various plant seeds, including wheat, celery, soybean, and others, and obtained fruitful achievements. Subsequently, the scientific team represented by Veselova analysed in detail the quality and performance of various crop seeds (soybean, barley, sunflower, etc.) using the delayed radiation of biophotons and found that there is a negative correlation between seed vigour and a delayed luminescence signal [10]. In recent decades, the research of biophoton technology has made tremendous development. A large number of experiments have proved that biophoton radiation is a common life phenomenon that is related to biological and physiological activities, the generation and synthetization of DNA, and other information exchange or energy transmission processes. The higher the level of an organism is, the greater intensity of the photon radiation that is emitted. The research applying of biophoton technology has been mainly conducted in medical fields, such as medical information diagnosis [11], cancer classification [12], analysis of brain activities [13], and others. In the cereal storage field, however, biophoton studies mainly focus on insect intrusion rather than on wheat mould. Duan et al. [14] have applied the permutation entropy algorithm to analyse the biophoton signals of wheat kernels and then use a BP network to test the experimental effects. Their proposed algorithm not only improves the detection rate by 10% but also saves the sample training time [14]. Regarding the detection process of insect intrusion, spontaneous biophoton emission, which is also known as ultra-weak luminescence (UWL), has been proven to be a sensitive index at reflecting the mould among wheat kernels. The main achievements in this paper are that we have measured the UWL of healthy and mouldy wheat kernels separately using biophotonic technology, calculated the approximate entropy and multiscale approximate entropy as the main classification parameters, and then, we sued the SVM to test the classification performance of newly established model.

## Materials and methods

### Materials

#### Wheat kernel samples

The experimental wheat kernels were offered by the Yuda grain barn, Zhumadian city, Henan Province in 2019. Some pretreatment, such as finding foreign materials and imperfect or damaged kernels, washing the kernels several times using distilled water, drying the samples to a certain degree of moisture using special equipment and so on, is very necessary. Subsequently, the wheat sample is divided into two parts: one is the healthy samples, and the other part is sent to the College of Biological Engineering to cultivate the mould in the sample with 50% Aspergillus flavus. Regarding the healthy wheat samples, we prepared 240 subsamples weighing 20.00±0.01 g. We use 120 subsamples as the training group (experimental group), and the remaining 120 subsamples form the testing group. Regarding the mouldy wheat samples, 120 subsamples are used as the training group, and the other 120 subsamples are used as the testing group. Meanwhile, protective measures should be taken during this process due to the strong poisonous of AFB1.

#### Equipment

The BPCL-2-ZL, manufactured by Beijing Jianxin Lituo Technology Co., Ltd., was used to measure the biophotons of healthy and mouldy wheat samples.

Fig. 2 shows the whole analysis system, which consists of three parts: ① a detection chamber, where the tested samples are input; ② a biophoton analyser, mainly including the photon counting and optical hi-voltage converter device; and ③ computer equipment, which displays results from the corresponding software on a monitor. The calculated average background noise of the instrument is 28 counts per second, The high voltage for the test is set as 1030 V, and the testing temperature is 25.0±0.5℃.

**Fig. 2.**
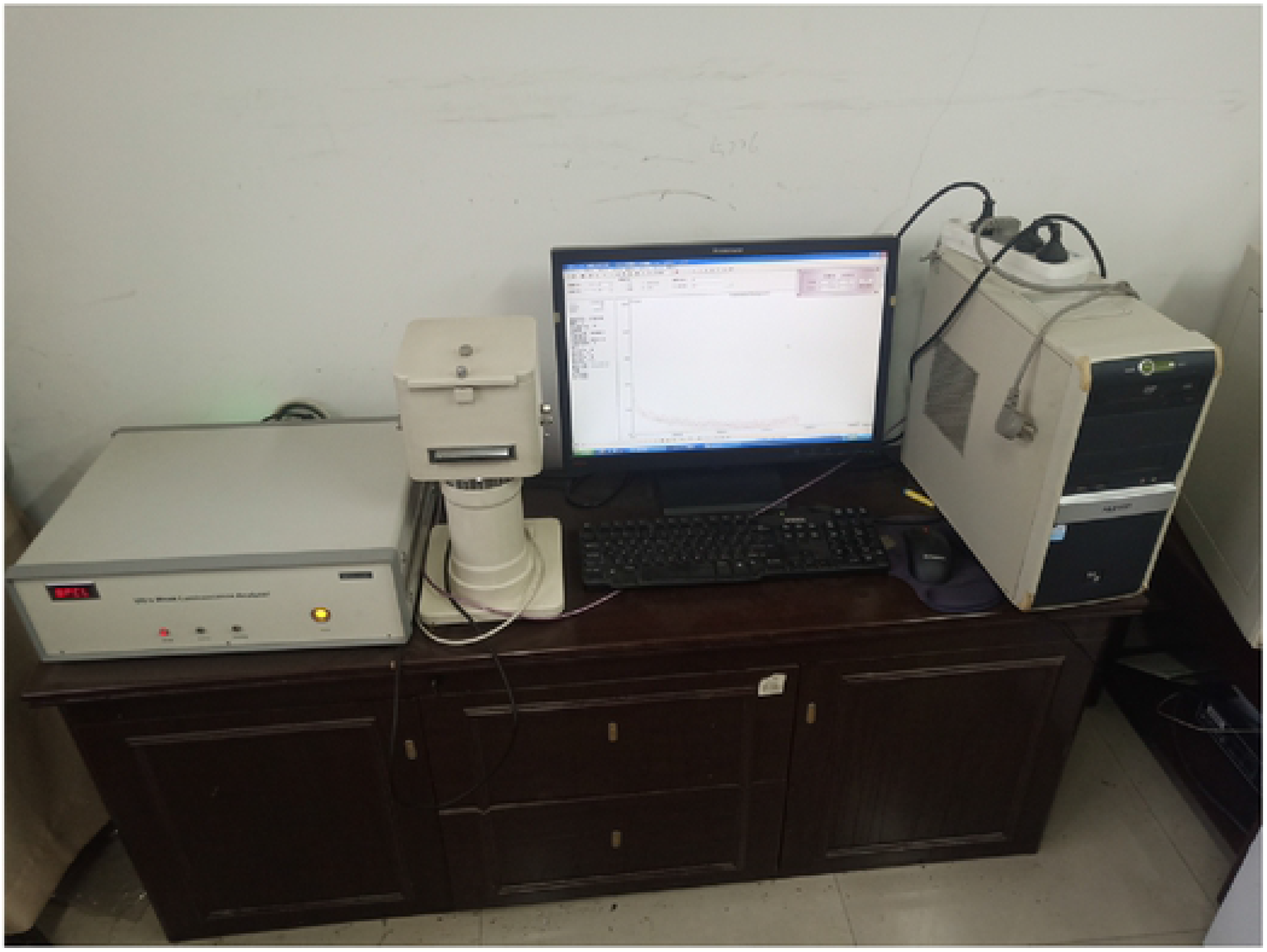
Instrumentation Used in the Experiment.

### Methods

The whole detection process consists of two parts. One part is selecting the right environmental parameters. Since the experimental result may be influenced by surrounding factors, all the experiments should be conducted under the same conditions to minimize the environmental influences, including the same environmental temperature (20±1℃), humidity (25±6%), and measuring time (8:00 am~18:30 pm). The other part is choosing suitable experimental parameters. Before testing, each sample was placed for 30 min in a dark space to decrease the interference from ambient parasitic light. Since the spontaneous biophotonic radiation of wheat kernels is not strong enough, the sampling interval is set to 10 s in order to collect ample numbers of biophotons. To better reflect the properties of the UWL signals of the two types of wheat samples, the total sampling time is extended over 15,000 s. Then, the UWL signals of the healthy and mouldy wheat kernel samples are measured separately.

## Results

### Biophotonic data analysis

One hundred and twenty groups of healthy and mouldy wheat samples were measured by the above processes. Owing to the nonlinear and random characteristics of the number of biophotons, we calculated the average numbers of photons for all the samples for both types, and the results are shown in Fig. 3. Table 1 shows their statistical characteristics, such as the mean, variance and standard deviation. As Table 1 shows, the statistical biophotonic characteristics of mouldy wheat are larger than those of healthy wheat. This difference occurs because the Aspergillus fungi that colonized the wheat kernels have much stronger metabolism and respiration. The large number of biophotons in mouldy wheat also provides a convincing explanation, which coincides with physiological regularity such that the higher the level of an organism is, the greater the intensity of the biophotons it emits.

**Fig. 3.**
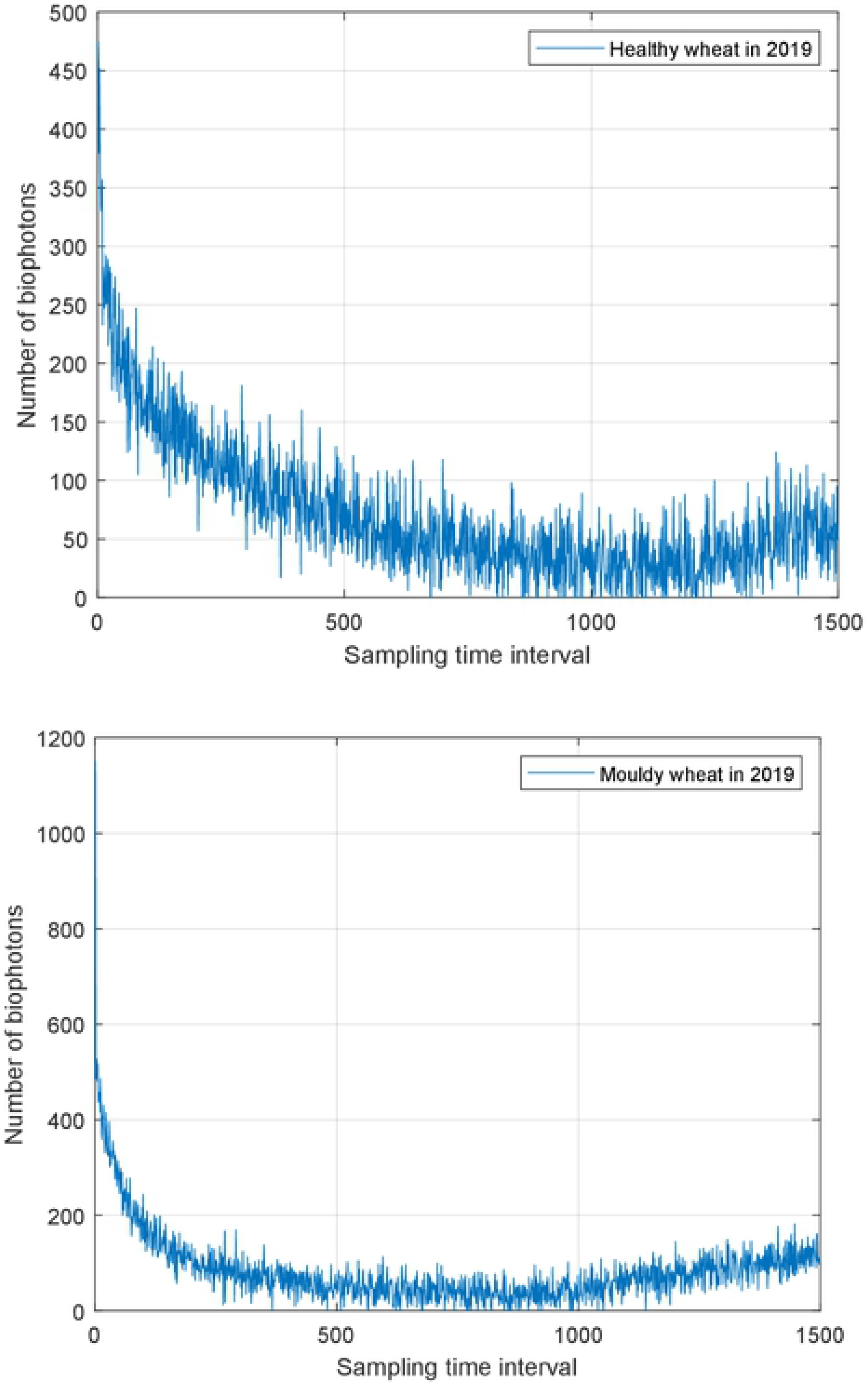
Average UWL Data of Healthy and Mouldy Wheat.

**Table 1.** UWL Data Statistical Characteristics of Two Types of Wheat.

|  | Mouldy wheat kernels in 2019 | Healthy wheat kernels in 2019 |
| --- | --- | --- |
| Mean | 84.22 | 71.11 |
| Variance | 5830.37 | 3533.45 |
| Standard deviation | 76.36 | 59.44 |

To effectively distinguish between healthy wheat and mouldy wheat based on UWL data, we will use the approximate entropy (ApEn) and multiscale approximate entropy (MApEn) algorithm, and then comparing their performances.

### Approximate entropy

The approximate entropy (ApEn) algorithm was proposed by the scholar Pincus to measure the characteristics of random series [15]. The more complex an initial time series is, the larger its corresponding ApEn. The ApEn is suitable for analysing the biophoton signals of wheat kernels because of its more robust performance. Two prominent advantages of the ApEn are its lower dependency on the length of the initial time series and strong resistance to the noise contained in the original data.

The complete computing process of the ApEn is [16]:

Divide the original series *X* ={*x*(*i*), *i* =1, 2,…, *N*} into an m-dimensional vector *u*(*i*) , which is shown as follows:

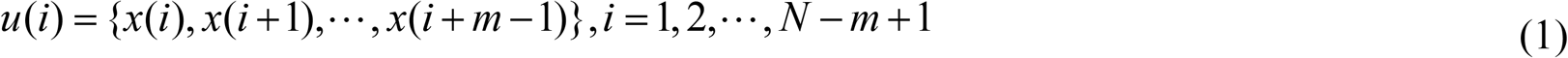

Here, *m* represents the dimension of the pattern vector, and *N* denotes the initial length of the time series.

1. Calculate the distance *d*[*u*(*i*), *u*(*j*)] between vector *u*(*i*) and vector *u*(*j*) using formula 2.

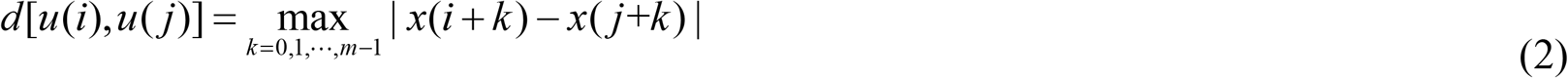
2. Count the numbers of *d*[*u*(*i*), *u*(*j*)]<*r*, where *r*, which is known as the similar tolerance threshold value, is a positive real number. Then, calculate the proportion between *d*[*u*(*i*), *u*(*j*)] < *r* and the total number of vectors, which is labelled as 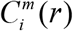 in equation 3.

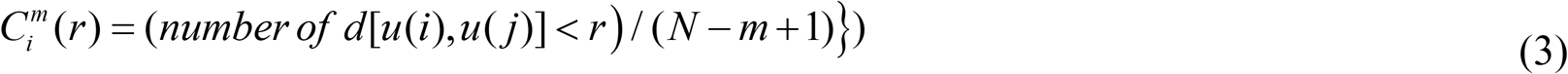

1. Calculate the logarithm of 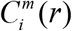, and then, obtain its mean using equation 4. Here, the mean is labelled as *H*^*m*^ (*r*).

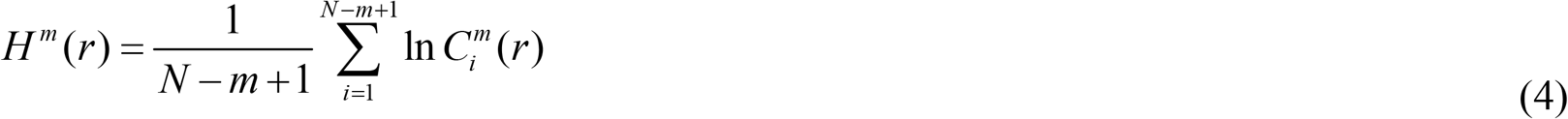
2. By increasing the dimension from *m* to *m* + 1 and repeating steps 2~4, *H*^*m*+1^ can be obtained.
3. The definition of the ApEn can be given as:

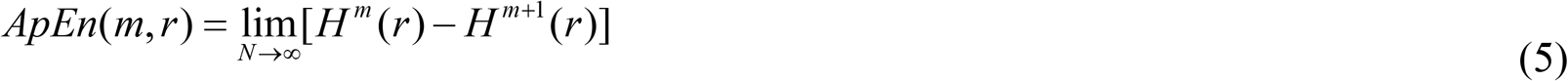 If *N* is finite, formula 5 is rewritten as:

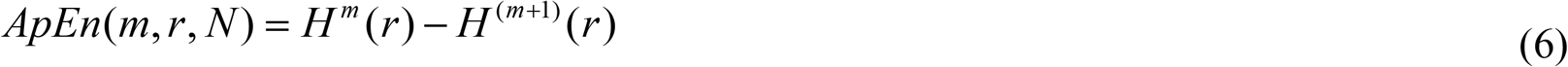

### Multiscale approximate entropy

To improve upon ApEn, the multiscale approximate entropy (MApEn) based on ApEn has been proposed to improve the robust and accuracy of model. Furthermore, the MApEn algorithm, overcomes the limitations of ApEn [17]. Interestingly, compared with only one feature obtained by ApEn, these MApEn values reflected by different scales are able to be used as a cluster of classification parameters for the subsequent SVM training model. The concrete steps of the MApEn algorithm are as follows [17]:

1. Assume the initial discrete series is *X* = {*x*(*i*), *i* = 1, 2,…, *N*}, and its length is N.
2. Construct a coarse time series {*z*(*τ*)} , where *τ* represents the scale factor, and then, the scaling time series can be expressed as:

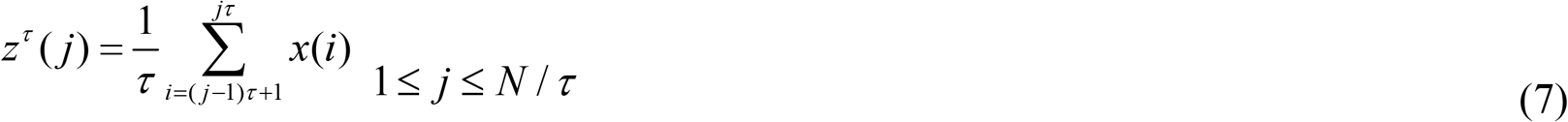

Equation 7 is the same as the original sequence provided that the scale factor *τ*=1 . Furthermore, each coarse-graining series can be regarded as evenly dividing the original series, and each segmentation length is *τ*.

By combining multiscales with the approximate entropy to generate MApEn, the MApEn algorithm is able to characterize the nonlinear information of series more effectively. Fig. 4 exhibits the detailed flowchart.

**Fig. 4.**
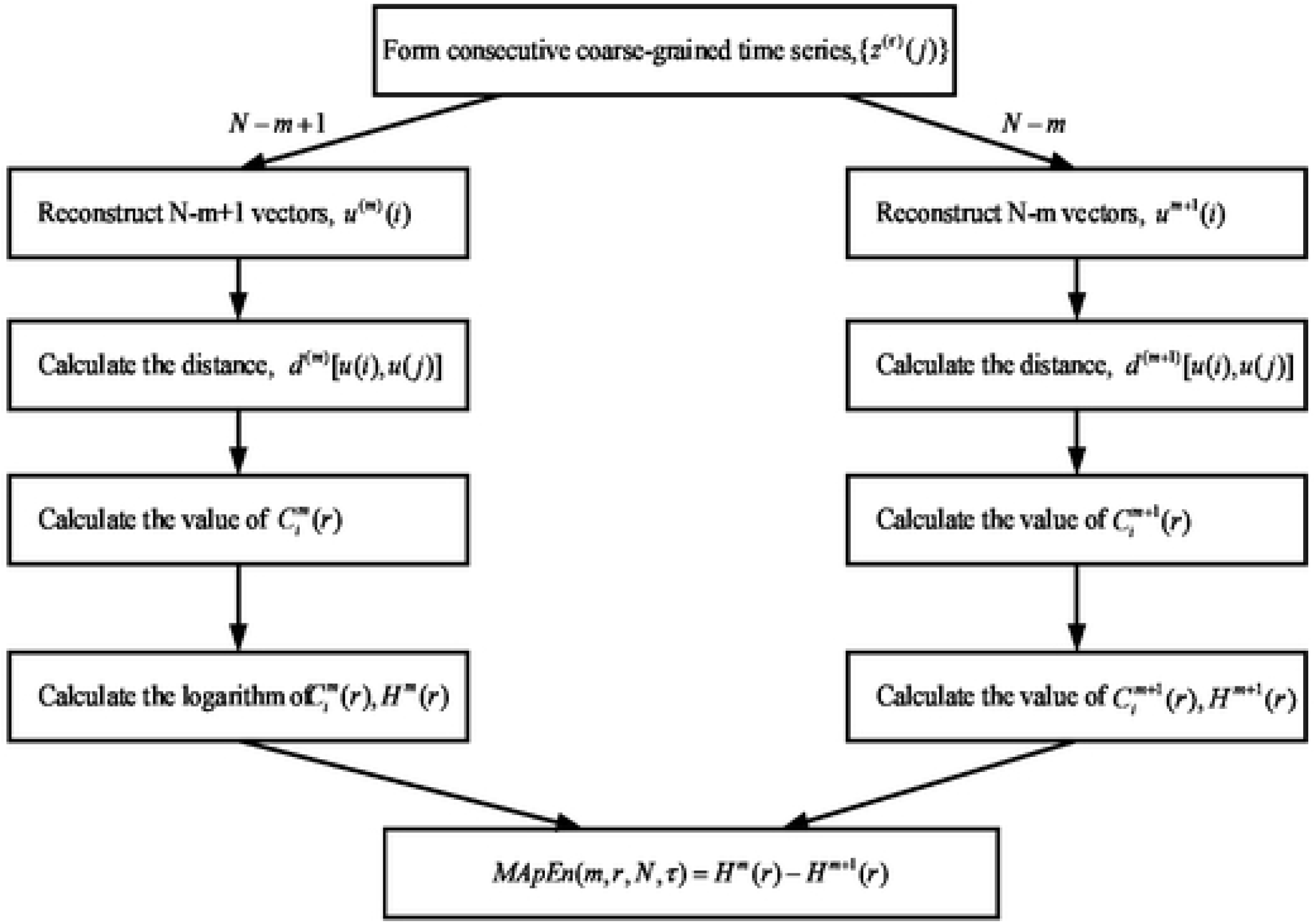
Flowchart of Multiscale Approximate Entropy Algorithm.

### MApEn algorithm and its performance

#### Fast ApEn algorithm and setting parameters

First, we can calculate the ApEn according to the abovementioned equations 2~6. There is plenty of redundant computing in some steps; however, it is time-consuming and cannot be used for real-time determination. Bo et al. [18] proposed a type of fast ApEn algorithm that can shorten the running time by nearly 5 times. The main steps are as follows:

First step: The distance matrix *D*(*N* × *N*) for the initial *N* points time sequence is calculated, and the element in the *i*^*th*^ row and *j*^*th*^ column can be denoted as *d*_*ij*_ . The rules for calculating *d*_*ij*_ are based on the following algorithm:

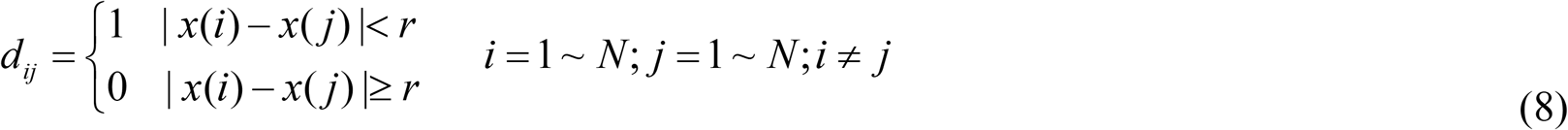

Second step: assuming the dimension of the pattern vector *m* = 2 , we can easily obtain the values of 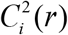 and 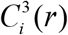 using equation 9.

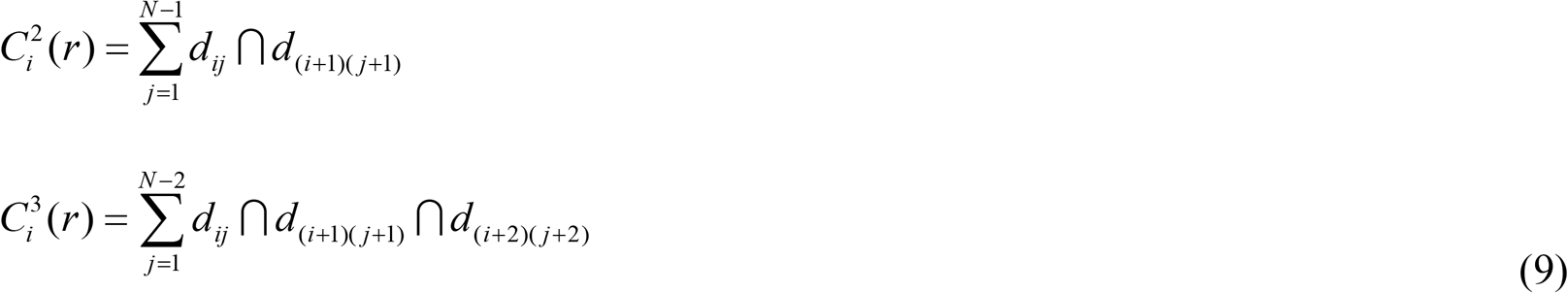

Third step: According to the values of 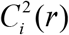 and 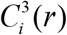, then we get *H*_2_(*r*) and *H*_3_(*r*).

Fourth step: The ApEn value can be calculated by equations 5~6.

Four parameters are involved in the MApEn algorithm: the length of the input signal *N* , the dimension of the pattern vector *m* , the similar tolerance threshold value *r* , and the time scale factor*τ*. For the ApEn algorithm, choosing the right parameters is of extreme importance to the algorithm.

After simulating several experiments, we finally select *N* = 1500, *m* = 2, *r* = 0.12 × *STD* as our experimental parameters, where STD represents the standard deviation of initial time series. The ApEn values of the UWL signal of the two types of wheat at different tolerance thresholds were simulated using Matlab 2018a, and the results are shown in Fig. 5. As shown in Fig. 5, the ApEn values of the two types of wheat vary depending on different tolerance thresholds *r* , Although the ApEn values of the two types of wheat are small, the differences between the healthy and mouldy wheat are obvious based on the ApEn values, where *r* varies from 0.1 to 0.19. In addition, another conclusion from the experimental results is that the smaller ApEn value of the mouldy wheat reflects that the activities of Aspergillus fungi are more regular and intensive than the healthy wheat itself, and thus, the value can be used as a classification feature to recognize mouldy wheat.

**Fig. 5.**
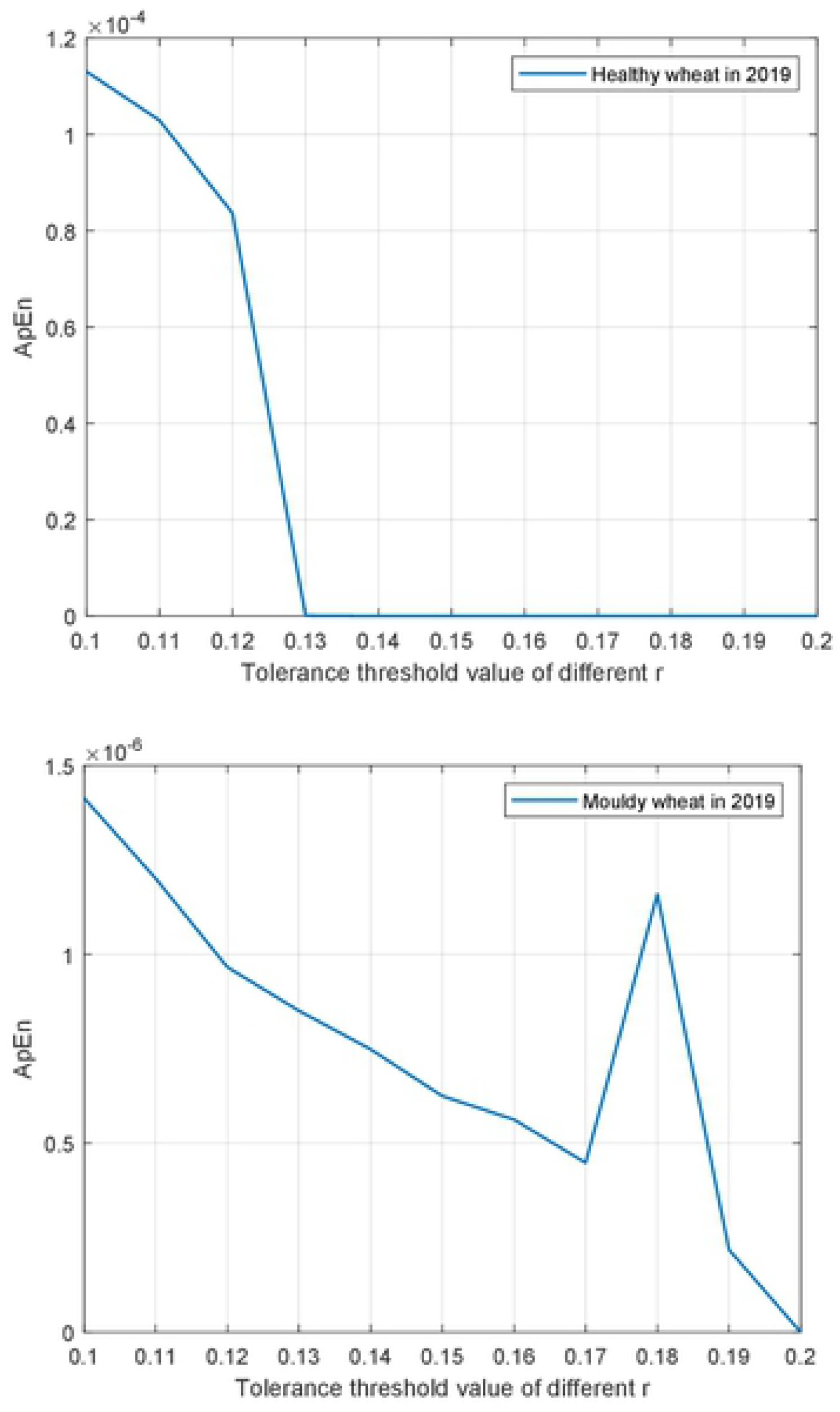
ApEn Values for Different Tolerance Thresholds of UWL Signals of Healthy and Mouldy Wheat.

## Discussions

### Performance analysis of the MApEn algorithm

The ApEn algorithm only offers one classification feature; therefore, in order to overcome this shortcoming and get more classification feature values, the MApEn algorithm is introduced in this paper. For ApEn algorithm, the parameters *N* = 1500, *m* = 2, *r* = 0.12 × *STD* are finally chosen and simulated via experiments. In addition to the parameters mentioned above, the scale factor *τ* is a decisive factor in the performance of the MApEn algorithm. Due to the limited length of the initial time series, *τ* is usually assigned a value from 2 to 10. The curve of the MApEn value at different scale factors is shown in Fig. 6.

**Fig. 6.**
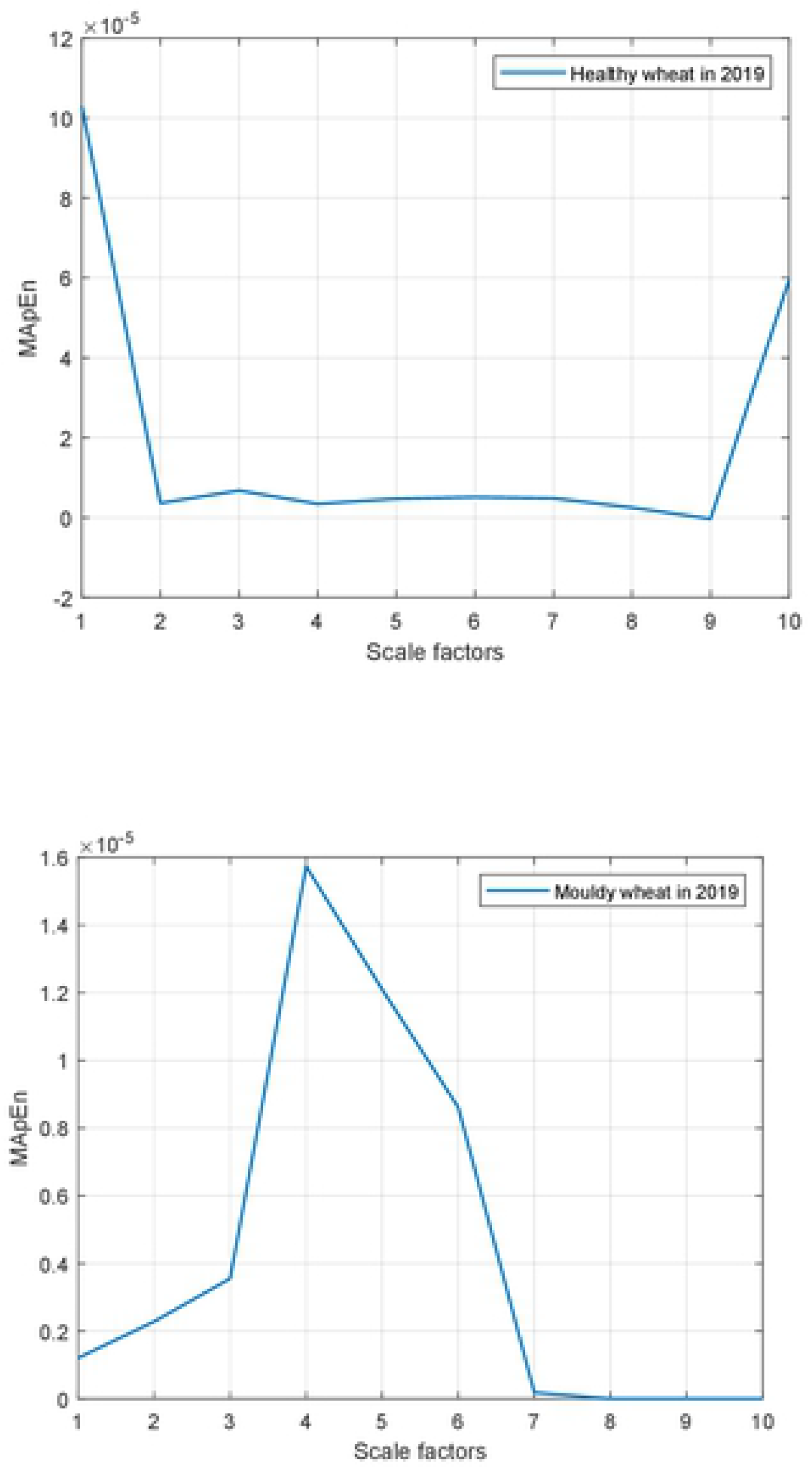
MApEn Values for Different Scale Factors of UWL Signals of Healthy and Mouldy Wheat.

Observing Fig. 6, the following conclusions can be achieved:

① The MApEn values of the UWL of the two types wheat sample shows an inverse trend;
② Compared with ApEn, the MApEn algorithm can offer several classification features that can be used at the same time under different scales rather than only one feature gained by ApEn algorithm.

### Bipartition classification and performance assessment by SVM

To solve the classification problem between healthy and mouldy wheat, the SVM is introduced in this work. The SVM, proposed by Cortes and Vapnik [16] in 1995, is a type of linear classifier based on classification boundaries. Computationally, the striking points of the SVM are how to choose the penalty and kernel parameters, and the kernel parameter impacts the nonlinear transformation of the input feature space from a lower-dimensional to a higher-dimensional space. In other words, this problem can be considered to be an optimization problem in which we seek to help the kernel function to find the optimal plane, by which we can conduct linearly separated classification based on a nonlinear transformation [19]. Although the training samples are not large enough, the SVM can achieve a good classification performance [20]. Currently, the SVM has become one of most widely used learning algorithm, and it has been applied in various fields [21,22].

Based on the SVM method and the purpose of the classification, the three parameters in Table 1 and the ApEn value act as the classification features. The UWL signals of total groups of the two types wheat kernels have been trained, and then, the abovementioned 120 healthy and mouldy wheat samples are separately used as the testing group. Adopting the SVM training model offered by Lin’s group from Taiwan University, the main parameters of the SVM are set as follows. The type of kernel function is a radial basis function, and the error value that terminates the iteration is 0.001. The ROC curve represents the classification result and is illustrated in Fig. 7, where the blue curve represents the classification performance of the MApEn algorithm, and the red curve represents the classification performance of the ApEn algorithm.

**Fig. 7.**
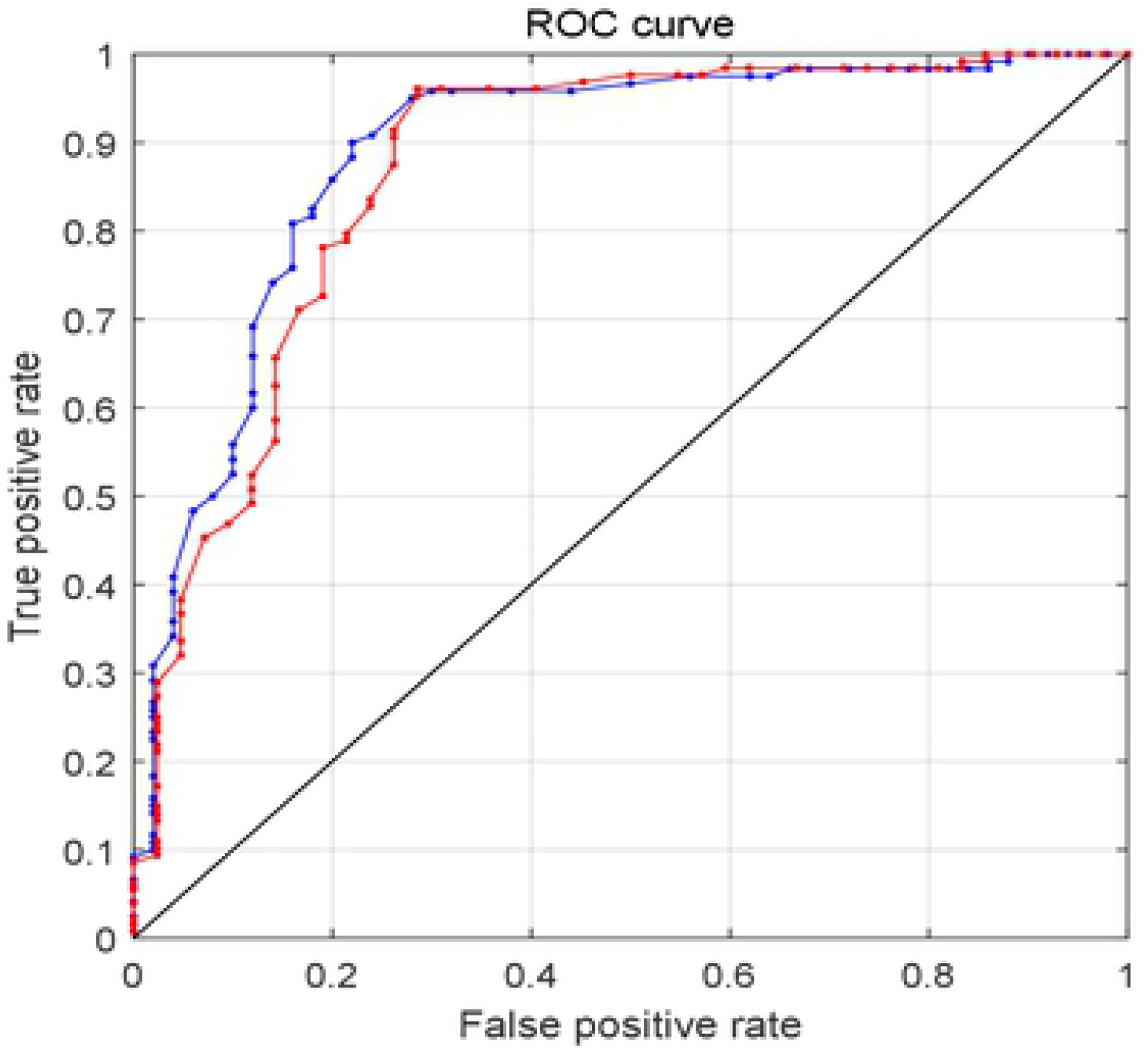
The ROC Curves of Two Classification Models.

From the ROC curves, Tables 2 and 3 can be calculated, where AUC, S.E., C.I., and PA represent the area under the curve, the standard error of the area, the confidence interval and the performance of the classifier, respectively. Comparing Table 2 with Table 3 shows that the classification accuracy rate based on MApEn has been improved obviously. In addition, the standard error decreases by introducing the MApEn algorithm. The experimental results validate that the MApEn values can act as a cluster of main classification features to recognize wheat kernels as healthy or mouldy.

**Table 2.** Classification Result Using ApEn as the Main Classification Feature.

| AUC | S.E. | 95% C.I. | PA |
| --- | --- | --- | --- |
| 0.8693 | 0.0272 | [0.8260 0.9226] | Good |

**Table 3.**
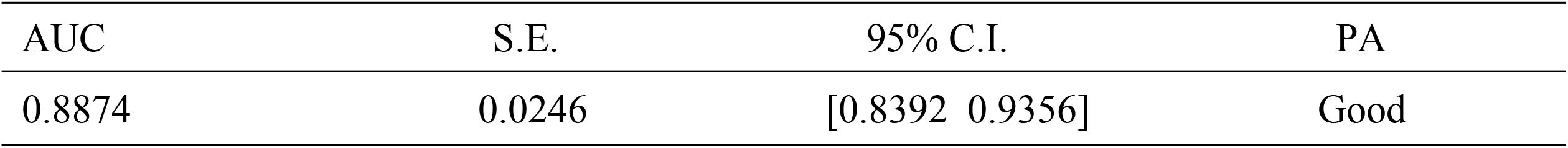
Classification Result Using MApEn as the Main Classification Features.

| AUC | S.E. | 95% C.I. | PA |
| --- | --- | --- | --- |
| 0.8874 | 0.0246 | [0.8392 0.9356] | Good |

## Conclusions

The UWL signals from different conditions of wheat kernels can reflect their inner physiological and pathological changes; therefore, it can be used as an environmentally friendly and nondestructive method to assess wheat quality. Since the UWL signal is so sensitive to environmental factors and the inner states of wheat kernels, further studies and experiments seeking to minimize these influences caused by these factors need to be conducted.

Multiscale approximate entropy is introduced to analyse the UWL signals in this paper. Subsequently, we have used an SVM to establish the classification model. The results of the simulations via an experiment show that the MApEn algorithm is efficient and effective at analysing random UWL signals. One main deficiency is that we only establish a binomial classification model in this work due to the limited experimental data, and the MApEn algorithm fails to exhibit its advantages. Furthermore, recognizing mouldy wheat kernels is a continuous process during their storage period; therefore, establishing a multiclassification model to classify and recognize the degree of mould is of extreme significance to help the operators to acquire accurate information about the degree of mould and make scientific choices, which requires further research to improve the precision of the established model.

## Acknowledgements

The authors are grateful for all the reviewers and the editor for their valuable suggestions and comments.

## References

1. USDA. Grain: world markets and trade. United States: Department of Agriculture Foreign Agricultural Service; 2018.

2. CHYXX. 2019. Available from: https://www.chyxx.com/industry/201910/793702.html.

3. Milani J. Ecological conditions affecting mycotoxin production in cereals: a review. Vet Med. 2013;58: 405–411.

4. Heshmati A, Zohrevand T, Khaneghah AM, Nejad ASM, Sant’Ana AS. Co-occurrence of aflatoxins and ochratoxin A in dried fruits in iran: dietary exposure risk assessment. Food Chem Toxicol. 2017;106: 202–208.

5. Blankson G, Mill-Robertson F. Aflatoxin contamination and exposure in processed cereal-based complementary foods for infants and young children in greater Accra, Ghana. Food Control. 2016;64: 212–217.

6. Mhiko TA. Determination of the causes and the effects of storage conditions on the quality of silo stored wheat (Triticum aestivum) in Zimbabwe. Nat Prod Bioprospecting. 2012;2: 21–28.

7. Gurwitsch A. Die natur des spezifischen erregers der zellteilung. Arch Mikrosk Anat Entwicklungsmechanik. 1923;100: 11–40.

8. Colli L, Facchini U, Guidotti G, Lonati RD, Orsenigo M, Sommariva O. Further measurements on the bioluminescence of the seedlings. Experientia. 1955;11: 479–481.

9. Popp FA, Gu Q, Li KH. Biophoton emission: experimental background and theoretical approaches. Mod Phys Lett B. 1994;8: 1269–1296.

10. Veselova T, Veselovsky V, Kozar V, Rubin A. Delayed luminescence of soybean seeds during swelling and accelerated ageing. Seed Sci Technol. 1988;16: 105–113.

11. Boschi F, Basso PR, Corridori I, Durando G, Sandri A, Segalla G, et al. Weak biophoton emission after laser surgery application in soft tissues: analysis of the optical features. J Biophotonics. 2019;12: e201800260.

12. Nirosha J, Murugan, Nicolas Rouleau, Lukasz M, Karbowski, Micheal A, et al. Biophotonic markers of malignancy: Discriminating cancers using wavelength-specific biophotons. Biochemistry and Biophysics Reports. 2018;13: 7–11.

13. Wang Z, Wang N, Li Z, Xiao F, Dai J. Human high intelligence is involved in spectral redshift of biophotonic activities in the brain. Proc Natl Acad Sci U S A. 2016;113: 8753–8758.

14. Duan S, Wang F, Zhang Y. Research on the biophoton emission of wheat kernels based on permutation entropy. Optik. 2019;178: 723–730.

15. Pincus SM. Approximate entropy as a measure of system complexity. Proc Natl Acad Sci U S A. 1991;88: 2297–2301.

16. Cortes C, Vapnik V. Support-vector networks. Mach Learn. 1995;20: 273–297.

17. Costa M, Goldberger AL, Peng CK. Multiscale entropy analysis of complex physiologic time series. Phys Rev Lett. 2002;89: 068102.

18. Bo H, Qingyu T, Fusheng Y, Tian-Xiang C. ApEn and cross-ApEn: property, fast algorithm and preliminary application to the study of EEG and cognition. Signal Process. 1999;15: 100–108.

19. Chapelle O, Vapnik V, Bousquet O, Mukherjee S. Choosing multiple parameters for support vector machines. Mach Learn. 2002;46: 131–159.

20. Zhang Y, Wu L. Classification of fruits using computer vision and a multiclass support vector machine. Sensors (Basel). 2012;12: 12489–12505.

21. Nagata F, Tokuno K, Mitarai K, Otsuka A, Ikeda T, Ochi H, et al. Defect detection method using deep convolutional neural network, support vector machine and template matching techniques. Artif Life Robot. 2019;24: 512–519.

22. Miyagi S, Sugiyama S, Kozawa K, Moritani S, Sakamoto SI, Sakai O. Classifying dysphagic swallowing sounds with support vector machines. Healthcare (Basel). 2020;8: 103.

